# Mu-opioid receptor system modulates responses to vocal bonding and distress signals in humans

**DOI:** 10.1101/2021.09.16.460660

**Authors:** Lihua Sun, Lasse Lukkarinen, Henry Karlsson, Jussi Hirvonen, Jari Tiihonen, Hannu Lauerma, Sophie Scott, Lauri Nummenmaa

## Abstract

Laughter is a contagious prosocial signal that conveys bonding motivation; adult crying conversely communicates desire for social proximity by signalling distress. Endogenous mu-opioid receptors (MORs) modulate sociability in humans and non-human primates. In this combined PET-fMRI study (n=17) we tested whether central MOR tone is associated with regional brain responses to social signals of laughter and crying sounds. MOR availability was measured with positron emission tomography using high-affinity agonist radioligand [^11^C]carfentanil. Haemodynamic responses to social laughter and crying sounds were measured using functional magnetic resonance imaging (fMRI). Social laughter evoked activation in the auditory cortex, insula, cingulate cortex, amygdala, primary and secondary somatosensory cortex, primary and secondary motor cortex; crying sounds led to more restricted activation in auditory cortex and nearby areas. MOR availability was negatively correlated with the haemodynamic responses to social laughter in primary and secondary somatosensory cortex, primary and secondary motor cortex, posterior insula, posterior cingulate cortex, precuneus, cuneus, temporal gyri, and lingual gyrus. For crying-evoked activations, MOR availability was negatively correlated with medial and lateral prefrontal haemodynamic responses. Altogether our findings highlight the role of MOR system in modulating acute brain responses to both positive and negative social signals.

## Introduction

Humans and nonhuman primates use numerous vocalizations for maintaining social bonds and proximity to their conspecifics. Laughter is a universally recognized positive social expression occurring frequently in human social interactions [1,2] and it is used for promoting social bonding [2,3]. Numerous other primates [4,5] and rodents [6] also use laughter-like vocalizations for conveying prosocial motivation. For example, in macaques and chimpanzees relaxed open-mouth vocalizations are associated with both play behaviour and pair formation [4,7]. Functional and acoustic properties of this kind of play signals are comparable in humans and other great apes, suggesting phylogenetic continuity on vocal communication of bonding motivation [8]. Crying is also used for signalling the need for social contact in humans and other mammals [9,10]. Unlike laughter it is evoked when distress or social distancing is experienced. Such distress cues engage the separation distress circuit in the mammalian brain that consequently modulate approach behaviour and social contact seeking [11]. Due to the centrality of human social attachment in well-being and mental health, it is imperative to understand the molecular systems that support processing of these distinct types of social attachment signals in the human brain.

There are numerous functional and molecular parallels in cerebral processing of social bonding signals conveyed by laughter and crying. Hearing adult laughter and crying activate amygdala, insula and auditory cortices [12–14], whereas hearing infant crying activates anterior insula, the pre-supplementary motor area and dorsomedial prefrontal cortex and the inferior frontal gyrus, as well as thalamus and cingulate cortices in adults [15]. At molecular level, human and animal studies converge in showing that the endogenous mu-opioid receptor (MOR) system modulating pleasurable and calm sensations [16] is an important mechanism for modulating social motivation [17]. *In vivo* molecular imaging studies in humans have shown that prosocial cues including social laughter trigger central endogenous opioid release [18], and that individual differences in MOR tone are associated with stable patterns of socioemotional behaviour such as childhood and adult romantic attachment styles [19,20]. Pharmacological studies in primates have further found that opioid receptor antagonists increase grooming and social behaviour [21,22]. Similarly to effect of social laughter, sustained sadness also induces endogenous opioid release in humans [23], while lowered endogenous MOR availability is associated with depressed mood [24]. Furthermore, MOR antagonist naltrexone amplifies negative feelings and subjective experience of pain when seeing others being hurt, suggesting the opioidergic modulation of empathy evoked by distress signals [25]. In line of this, animal studies also suggest that opioid agonists alleviate and antagonists potentiate separation distress as quantified by crying-like distress vocalizations [26], suggesting opioidergic contribution in processing distress signals. Against this background, it may be expected that individual variation in MOR availability could be associated with acute regional responses to social signals of laughter and crying. However, *in vivo* evidence regarding the role of the MOR system in acute responses to social signals conveyed by laughter and crying is currently elusive.

### The current study

Here we used fusion imaging with PET and fMRI to delineate the functional and molecular brain systems involved in processing of social signals conveyed by laughter and crying. We measured haemodynamic responses to social laughter and crying sounds using fMRI while baseline MOR availability was quantified with PET using high-affinity agonist radioligand [^11^C]carfentanil. We then predicted haemodynamic responses to laughter and crying with regional MOR availabilities. We show that MOR availability is linked with haemodynamic responses to both laughter and crying, but that the spatial layout of these MOR-BOLD interactions is distinct for the different vocalization types.

## Methods

### Subjects

Seventeen healthy males (Age 29.2 ± 7.9; BMI 25.0 ± 2.2) volunteered for the study. The study was approved by the ethics committee of the hospital district of South-Western Finland. The study was conducted according to the declaration of Helsinki and all subjects provided written consents for participating the study.

### PET data acquisition and preprocessing

PET data were acquired using a GE Healthcare Discovery 690 PET/CT scanner. PET images were preprocessed using the automated PET data processing pipeline Magia [27] (https://github.com/tkkarjal/magia) running on MATLAB (The MathWorks, Inc., Natick, Massachusetts, United States). Radiotracer binding was quantified using non-displaceable binding potential (BP_ND_), calculated as the ratio of specific binding to non-displaceable binding in the tissue [28]. This outcome measure is not confounded with differences in peripheral distribution or radiotracer metabolism. BP_ND_ is traditionally interpreted by target molecule density (Bmax), although [^11^C]carfentanil is also sensitive to endogenous neurotransmitter release. Accordingly, the BP_ND_ for the tracer should be interpreted as density of the receptors unoccupied by endogenous ligand (i.e., receptor availability). Binding potential was calculated by applying basis function method [29] for each voxel using the simplified reference tissue model [30], with occipital cortex serving as the reference region [31]. The parametric images were spatially normalized to MNI-space via segmentation and normalization of T1-weighted anatomical images, and finally smoothed with an 8-mm FWHM Gaussian kernel.

### fMRI data acquisition and analysis

#### Experimental design and stimuli

In the vocal expression fMRI task, the subjects listened to short laughter and crying vocalizations, or control stimuli that were generated by time-domain scrambling of the original sounds. The original stimuli have been validated and described in detail in [32]. The experiment was run using a blocked design. In each 16.5s block, five 2.5s stimuli from one category (i.e., laughter, crying sounds, scrambled laughter, or scrambled crying sounds) were played with a 1s silent period between stimuli (**Fig 1A**). The blocks were interspersed with rest blocks lasting for 4-7s. To keep participants focused on the task, an animal sound (vocalization of an alpaca for 3s) was presented randomly with 50% chance during the rest blocks. The subjects were instructed to press the response button whenever they heard the alpaca, and the behavioural responses were inspected to guarantee the focus of attention. A total of 32 blocks (8 blocks per stimulus type) were run in randomized order.

**Figure 1.**
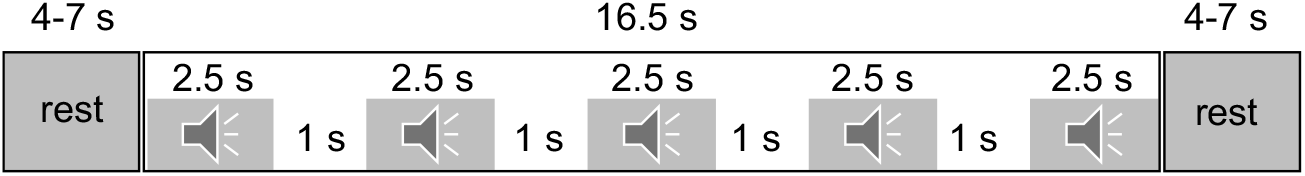
Experimental block design.

#### fMRI data acquisition and preprocessing

Phillips Ingenuity TF PET/MR 3T whole-body scanner was used for collecting the MRI data. Structural brain images with resolution of 1 mm^3^ were acquired using a T1-weighted sequence (TR 9.8 ms, TE 4.6 ms, flip angle 7°, 250 mm FOV, 256 × 256 reconstruction matrix). Brain structural abnormalities were screened by a radiologist (JH). Functional MRI data were acquired using a T2*-weighted echo-planar imaging sequence (TR = 2600 ms, TE = 30 ms, 75° flip angle, 240 mm FOV, 80 × 80 reconstruction matrix, 62.5 kHz bandwidth, 3.0 mm slice thickness, 45 interleaved slices acquired in ascending order without gaps). A total of 290 functional volumes were acquired.

MRI data were processed using the fMRIPrep 1.3.0.2 [33]. Structural T1 images were processed following steps: correction for intensity non-uniformity, skull-stripping, brain surface reconstruction, spatial normalization to the ICBM 152 Nonlinear Asymmetrical template version 2009c [34] using nonlinear registration with antsRegistration (ANTs 2.2.0) and brain tissue segmentation. Functional MRI data were processed as follows: co-registration to the T1 reference image, slice-time correction, spatial smoothing with a 6mm Gaussian kernel, automatic removal of motion artifacts using ICA-AROMA [35] and resampling to the MNI152NLin2009cAsym standard space. Image quality was inspected visually for the whole-brain field of view coverage, proper alignment to the anatomical image. Signal artifacts were assessed via the visual reports of fMRIPrep. All functional data were thereafter included in the current study.

#### fMRI data analysis

The fMRI data were analysed in SPM12 (Wellcome Trust Centre for Imaging, London, UK, (http://www.fil.ion.ucl.ac.uk/spm). The whole-brain random effects model was applied using a two-stage process with separate first and second levels. For each subject, GLM was used to predict regional effects of task parameters on BOLD indices of activation. Contrast images were generated for laughter or crying sound versus corresponding scrambled sounds and subjected to second-level analyses. Statistical threshold was set at p < 0.05, FDR corrected at cluster level.

### PET-fMRI fusion analysis

#### Region of interest definition

Seventeen ROIs were selected based on i) their role in emotional processing and high ii) MOR expression [16,36,37]. These ROIs include the frontal pole (FP), insula, orbitofrontal cortex (OFC), anterior cingulate cortex (ACC), and posterior cingulate cortex (PCC), and precuneus (PreCu), amygdala, thalamus, ventral striatum, caudate, putamen, hippocampus (HC) defined by the AAL atlas [38]. We also included the subregions of motor area, given their important role processing social stimuli [39,40]; they were was parcellated in the Juelich Atlas with masks generated using the SPM Anatomy toolbox [41]. These subregions include the primary motor cortex (M1) corresponding to Brodmann areas (BA) 4a and 4b, the supplementary motor area (M2) corresponding to BA6 [42] the primary somatosensory cortex (S1) including BA3a, BA3b, BA1 and BA2 [43,44], and the secondary somatosensory cortex (S2) including parietal operculum 1-4 [45]. Finally, the auditory cortex was defined using the Juelich Atlas combining TE 1.0, TE1.1 and TE 1.2 [46] was included in the ROI set. Mean regional MOR availabilities were extracted for each ROI.

#### Fusion analysis

Two different approaches were used for fusion analysis. In the full-volume approach, voxel-wise BOLD responses to laughter and crying were predicted with ROI-wise [^11^C]carfentanil availabilities (i.e. separately for each ROI) using linear regression analysis. Statistical threshold was set at p < 0.05, FDR corrected at cluster level. In a complementary methodological approach, we also extracted subject-wise BOLD responses to laughter and crying in the 17 ROIs described above. Subsequently, MOR availabilities in these ROIs were correlated (Pearson) with the corresponding regional BOLD responses to characterize the regional interactions between MOR and BOLD responses to laughter and crying. Correlation analysis was conducted with R statistical software (version 3.6.3).

## Results

**Figure 2** shows the mean MOR distribution in the subjects. MORs are widely distributed across the frontal, temporal, parietal and subcortical brain regions.

**Figure 2.**
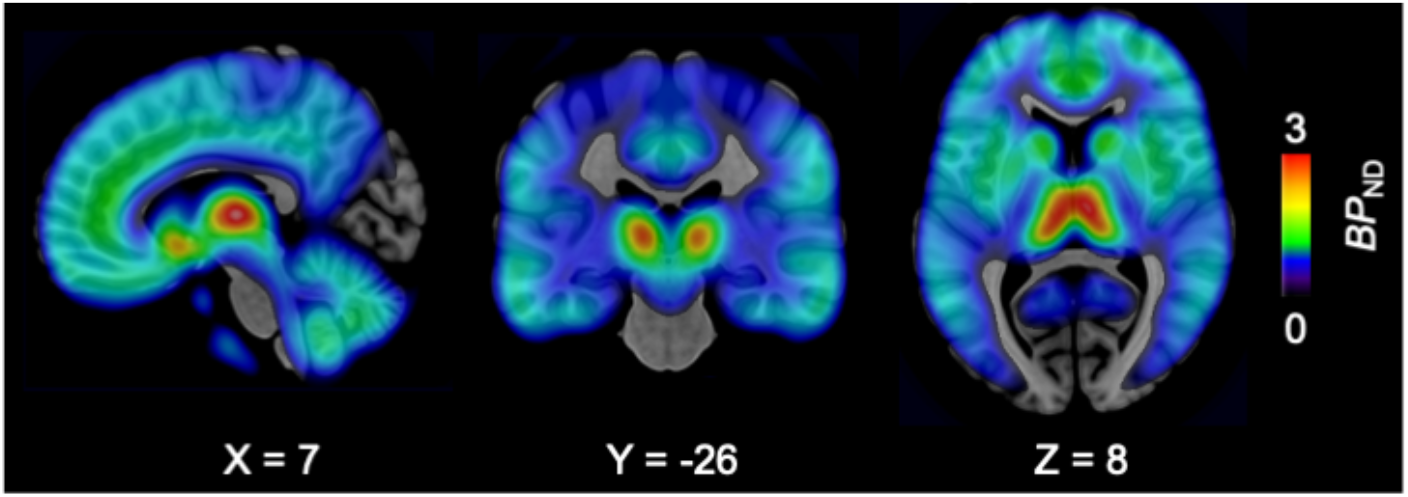
Mean distribution of MORs in the subjects (*n*=17).

Laughter versus scrambled laughter elicited activation in primary and secondary auditory cortices and adjacent temporal regions, anterior (ACC) and posterior cingulate cortex (PCC), primary (S1) and secondary somatosensory (S2) cortex, primary (M1) and secondary (M2) motor cortex, medial frontal cortex, insula, amygdala, hippocampus, striatum and thalamus (**Fig 3A**). Crying sounds versus scrambled crying sounds activated only the primary and secondary auditory cortices and adjacent superior and middle temporal regions (**Fig 3B**). Direct contrast between laughter and crying showed significantly stronger activations in regions including M1, S2, thalamus, ACC and PCC, whereas the opposite contrast did not reveal any significant activations (**Fig 3C**).

**Figure 3.**
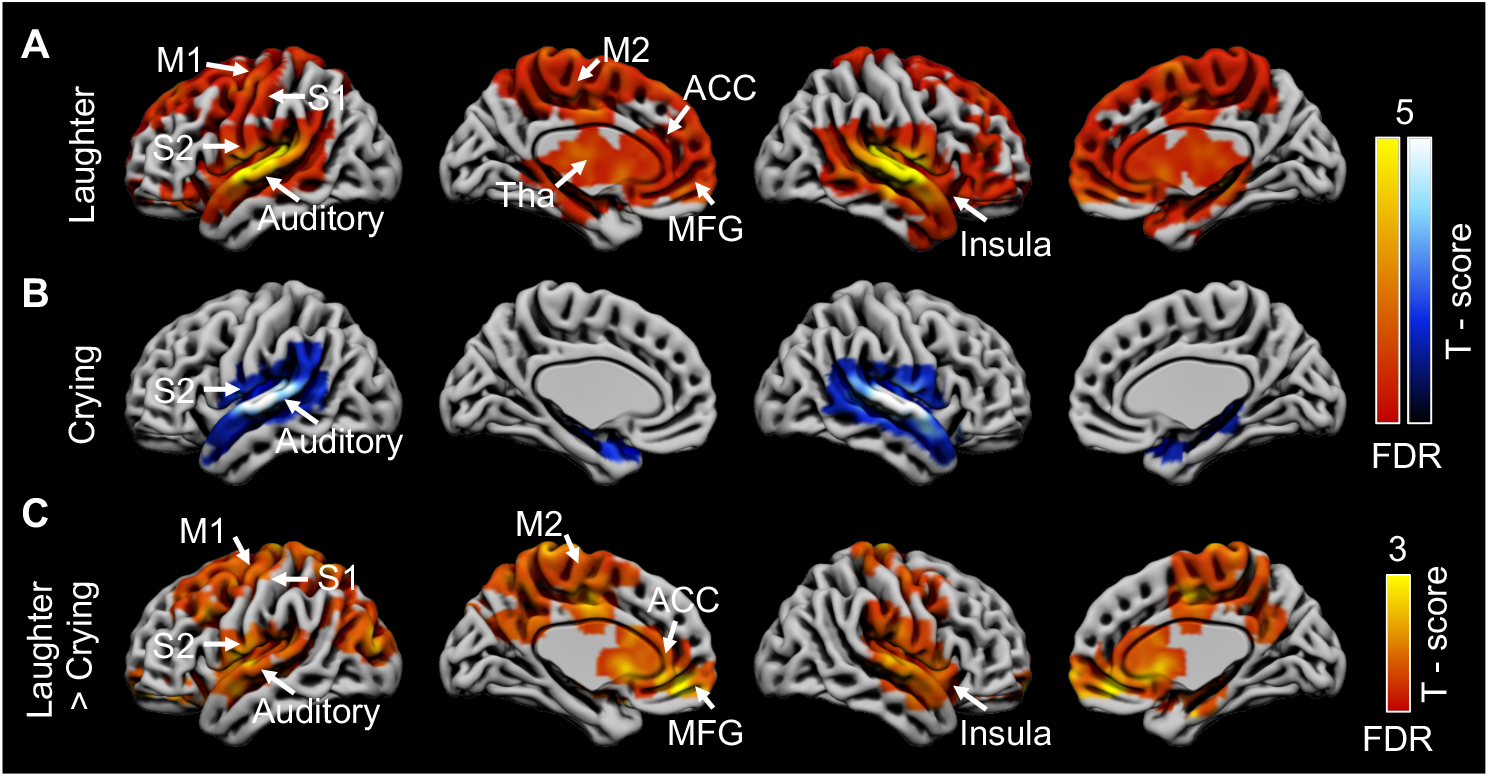
Haemodynamic responses to laughter and crying. **A.** BOLD responses to laughter vs. scrambled laughter (hot colours). **B.** BOLD responses to crying vs. scrambled crying sound (cool colours). **C.** Increased brain activation to laughter versus crying sounds in the interaction contrast (laughter – scrambled laughter) > (crying – scrambled crying). Results are FDR-thresholded at p < 0.05. M1 = primary motor cortex, M2 = secondary motor cortex, S1 = primary somatosensory cortex, S2 = secondary somatosensory cortex, ACC = anterior cingulate cortex, MFG = medial frontal gyrus.

### Fusion analysis

Next, we used regional MOR availabilities to predict BOLD responses to laughter and crying. In general, the associations were negative, but the spatial distribution of the effects was markedly different for laughter and crying. **Figure 4** shows cumulative maps where voxel intensities indicate the number of ROIs (out of 17) whose [11C]carfentanil BPND was correlated (p < 0.05, FDR corrected) with BOLD responses to laughter and crying in each voxel. For laughter, most consistent effects spanned the posterior cortical areas including the M1, M2, S1, S2, posterior insula, PCC, inferior, middle and superior temporal gyri, precuneus, cuneus, lingual gyrus. For crying, the most consistent effects were found for frontal cortical areas (including the inferior, middle, superior and medial frontal gyri) and anterior insula. No positive correlations were found for laughter. For crying sound only limited number of ROIs showed positive correlations with BOLD signals (**supplementary Figure S1**), and these were limited in scope. Region-wise fusion analysis maps are shown in **supplementary Figure S2**.

**Figure 4.**
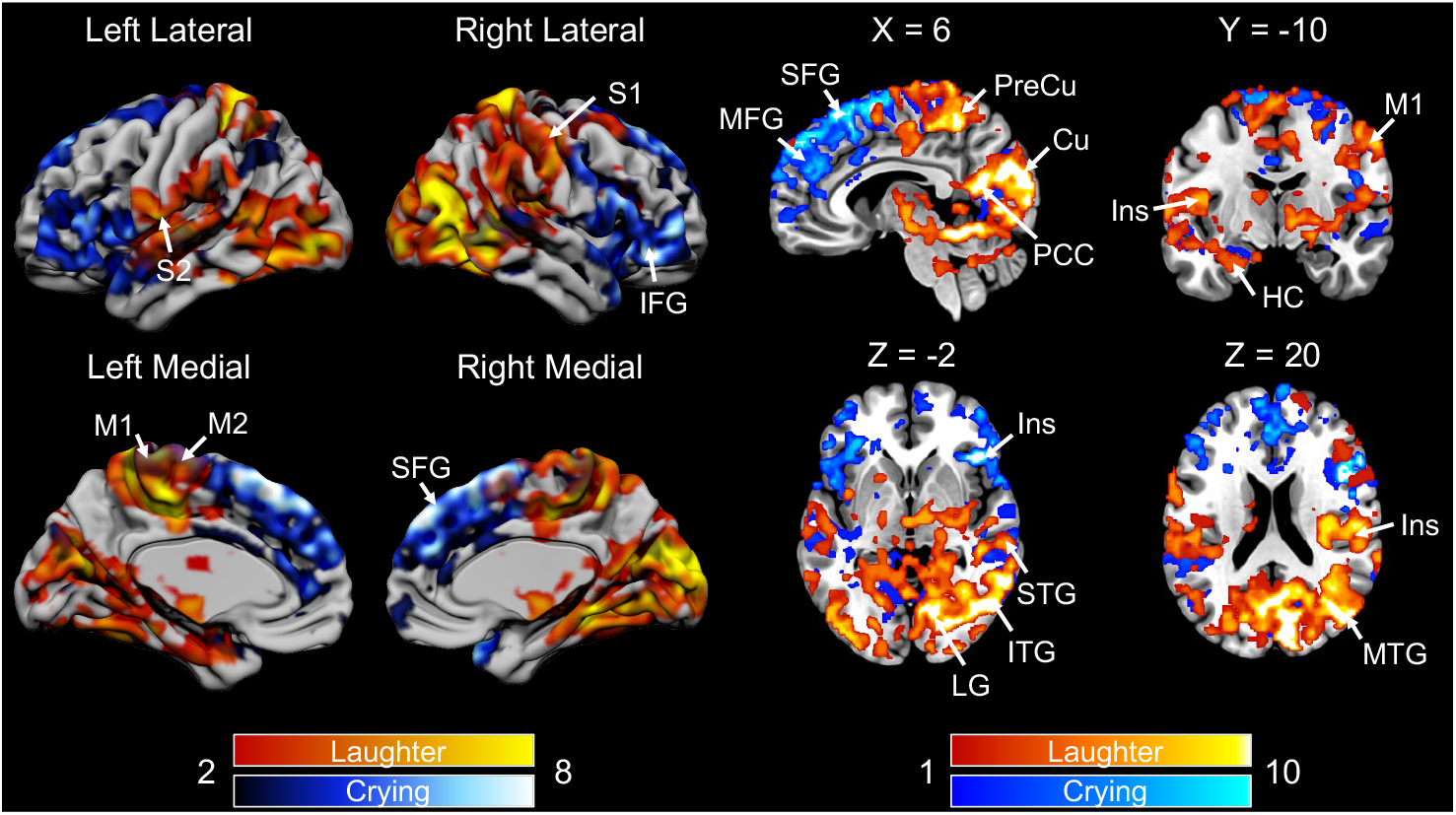
Cumulative maps indicating the number of ROIs (out of 17) whose [^11^C]carfentanil BP_ND_ was correlated with BOLD responses to laughter (hot colours) and crying (cool colours) in each area. M1 = primary motor cortex, M2 = secondary motor cortex, S1 = primary somatosensory cortex, S2 = secondary somatosensory cortex, MFG = medial frontal gyrus, SFG = superior frontal gyrus, PreCu = precuneus, Cu = cuneus, PCC = posterior cingulate cortex, Ins = insula, HC = hippocampus, STG = superior temporal gyrus, ITG = inferior temporal gyrus, LG = lingual gyrus.

### ROI-level correlations between MOR and BOLD responses

For laughter, only secondary somatosensory cortex (S2) showed a correlation between MOR availability and BOLD responses (**Fig 5A**). For crying, no regional correlations were found. In addition to within-region correlations, S2 MOR availability was correlated with BOLD response to laughter in auditory cortex. For crying, there were significant correlations between MOR availability in M2 and BOLD signals in amygdala and the auditory cortex (**Fig 5B**), and these correlations were absent for social laughter. Expanded plot of inter-ROI correlations is shown **supplementary Figure S3-4**.

**Figure 5.**
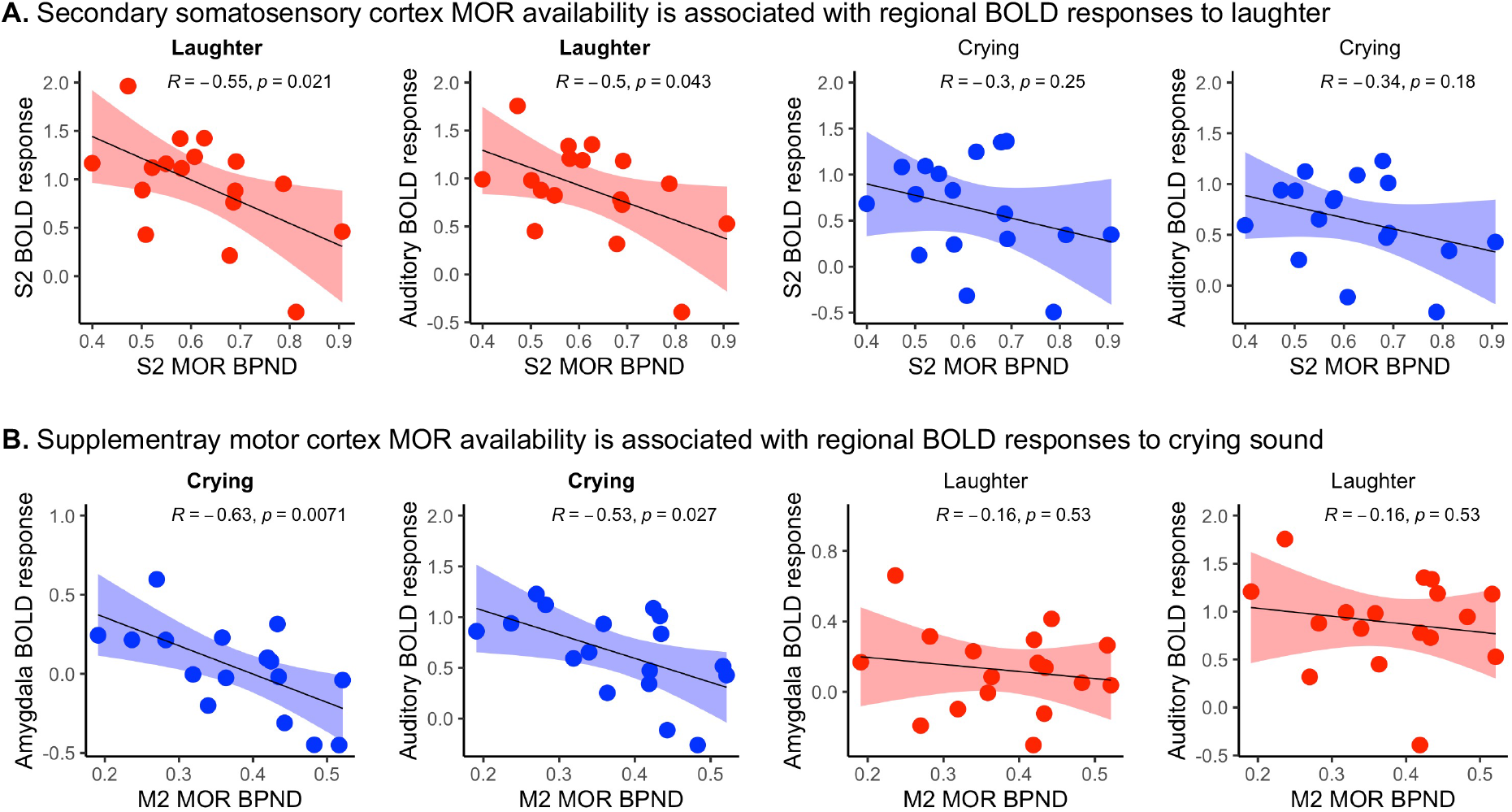
ROI-level correlations between MOR availability and BOLD responses to laughter (**A**) or crying sound (**B**). Shaded area shows 95% CI. M2 = secondary motor cortex, S2 = secondary somatosensory cortex.

## Discussion

Our main finding was that individual differences in cerebral MOR availability are associated with functional BOLD responses to both laughter and crying sound, yet the MOR-dependent responses to laughter and crying have distinct topographic patterns. MOR availability was associated with haemodynamic responses to laughter in somatosensory and motor cortices, posterior insula and temporal gyri, whilst the corresponding effects for crying were focused in the medial and lateral frontal cortex and anterior insula. These data extend the prior work on MOR-dependent individual differences in trait-level sociability by showing that the MOR system governs human sociability also via modulating acute processing of both prosocial and distress signals.

### MORs modulate responses to prosocial signals

The BOLD-fMRI analysis revealed that while both laughter and crying vocalizations reliably activated the auditory cortices, laughter also increased activity throughout the cingulate and somatosensory and motor cortices. Furthermore, laughter versus crying evoked significantly stronger limbic and paralimbic activation patters. Laughter is a pleasant prosocial signal that is highly contagious [40], and prior studies have consistently indicated that laughter also induces activation of the motor and premotor areas [47]. This kind of “mirroring” of social laughter may serve social bonding purposes, as it allows laughter to effectively spread across large crowds [3]. At neuromolecular level, social laughter evokes endogenous opioid release – an important neurochemical response promoting social bonding [48]. Presumably, the laughter-induced opioid release and concomitant pleasurable and relaxing sensations act a as a safety signal, promoting future social engagement with the current interaction partners. The present data imply that baseline MOR availability could be a neurochemical proxy for individual differences in responsiveness to prosocial cues, as the haemodynamic responses to laughter vocalizations depended on individual-specific MOR levels. These effects were consistently observed in the somatosensory and motor cortices. This finding accords with prior work on the role of the MOR system in human social bonding which have shown that individual differences in MOR tone are associated with trait-like differences in social bonding motivation [19,20]. Further, one human PET study has found that social laughter increases opioid release in the thalamus and insula, and that endogenous MOR tone positively predicted the occurrence of laughter during social interaction [18]. Moreover, social laughter is associated with increased pain threshold - an indirect assay of endogenous opioid release [49], and pharmacological studies in nonhuman primates suggest that opioid agonists and antagonists have a causal role in modulating social bonding behaviour [22,50]. Here we extended these findings by showing that MOR tone also links with acute functional responses to perceiving others’ social bonding cues, i.e. the higher opioid tone an individual had, the weaker the haemodynamic responses to laughter.

### MORs and distress signal processing

Previous PET studies have established that MOR system modulates responses to affiliative social cues such as laughter and social touching [18,48]. We extended these data by showing that MORs also modulated processing of distress vocalizations, possibly reflecting a MOR-mediated empathetic response. This accords with prior pharmacological studies showing that blocking the MOR signalling with naltrexone increases attention to both angry and happy facial expressions [51], thus implicating MOR-modulated vigilance towards both positive and negative social signals. Molecular imaging studies in humans have found that sustained sadness induces endogenous opioid release in humans [23], and social rejection may trigger transient changes in endogenous opioid peptide release [52]. Furthermore, opioid-mediated placebo analgesia reduces empathetic concerns and activity in the brain’s empathy circuit when seeing others in pain. Conversely, the MOR blocker naltrexone increases negative feelings and subjective experience of pain when seeing others being hurt [25]. Finally, one previous PET-fMRI study found that striatal MOR availability is negatively associated with haemodynamic responses in thalamus, postcentral gyrus and insula during pain observation [37]. Although this study used naturalistic and uncontrolled video stimuli, these results support the general role of MORs in modulating responses to distress cues.

While MORs modulated responses to prosocial (laughter) vocalizations primarily in the somatomotor and parietotemporal areas, the MOR-dependent responses to distress vocalizations were found in the prefrontal lobe. This was most prominently observed in the medial and prefrontal cortex, which is well known for its role in for social inference and decision making [53], and structural imaging studies has shown that frontocortical volumes are associated with the brevity of human social networks [54,55]. The present studies raise the possibility that MOR-mediated responses to others’ distress in the frontal cortex could be a putative mechanism leading to helping those who are in distress and concomitant strengthening of social bonds, highlighting the MOR-dependent modulation of social motivation [56].

We observed, in general, a negative association between MOR availability and haemodynamic responses to laughter and crying. This accords with prior fusion imaging studies showing that high MOR tone may downregulate acute haemodynamic responses to both distressing and arousing socioemotional events [37,57]. Yet, studies linking opioid receptor signalling with trait measures of sociability have typically observed a positive association between sociability and MORs [19,20]. One way to reconcile these lines of evidence is that while heightened MOR availability is linked with increased trait-level sociability, it is simultaneously associated with weaker responses to transient bonding and distress signals, and opioid system may thus modulate social perception and behaviour at multiple timescales. Therefore, it is possible that individuals with higher MOR availability and thus stronger prosocial motivation have more efficient processing of social signals (as indexed by lower BOLD response amplitudes) due to their repeated exposure to socioemotional cues. However, the present cross-sectional study does not allow us to test this hypothesis directly.

## Limitations

Our data are cross-sectional, and therefore we cannot conclude whether the links between MOR availability and responses to bonding/distress cues reflect i) genetically determined individual differences in MOR availability [58] contributing to differential patterns of social responsiveness or ii) downregulation of MOR neurotransmission resulting from different social environments and social interaction patterns. We also only scanned male participants and our results may not generalize to females. Our sample size was limited due to the complex PET-fMRI instrumentation; however the MOR density estimates based on [11C]carfentanil PET are reliable even in small samples such as the current one [59]. Finally, a single baseline scan cannot determine the exact proportions for causal factors leading to altered receptor availability which may be affected by changes in receptor density, affinity or endogenous ligand binding [60].

## Conclusions

We conclude that the central MOR system modulates processing of both prosocial and distress vocalizations with different regional contributions. This significantly extends the prior human work that has so far confirmed the contribution of MOR system to prosocial cues. The present results also highlight that baseline MOR tone predicts acute neural responses to both affiliative and distress cues, implying that MOR expression could underlie individual differences in social and affiliative behaviour. Because social attachment patterns are established over repeated exposures to others social signals, individual differences in MOR tone could explain why some individuals are more sensitive for responding to social signals and consequently more likely to establish social bonds. Taken together, the MOR system is broadly linked with processing of multiple aspects of human social signals, and it may contribute to modulation of social closeness both when others are in distress and seeking for social contact for enjoyment.

## Acknowledgements and funding statement

The study was supported by the Academy of Finland (grants numbers 294897 and 332225, to L.N.), Valon Vuoksi Foundation (grants to L.S. and L.L.) and Turku Collegium for Science and Medicine, University of Turku (to L.S.). We thank the imaging team, especially the research nurses, at the Turku PET Centre for facilitating the data collection.

## Author contributions

Conceptualization: L.S., J.T., H.L., S.S., L.N.; Data curation: L.S., L.L., H.K., J.H., H.L., L.N.; Formal analysis: L.S.; Supervision: L.N.; Visualization: L.S., L.N.; Writing – original draft: L.S., L.N.; Writing – review & editing: L.L., H.K., J.H., H.L., S.S.

