## Supplementary Data for "Mu-opioid receptor system modulates responses to vocal bonding and distress signals in humans"

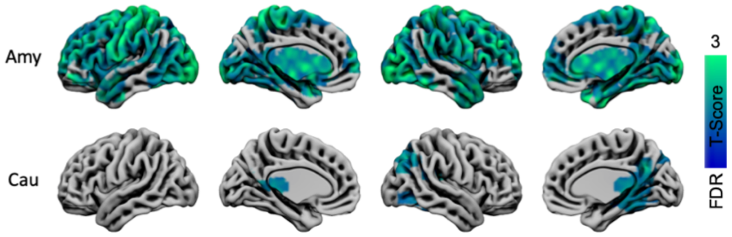


**Figure S1.** Positive correlations with regional MOR and brain BOLD response to crying sound. No positive correlations were found for laughter. Results are FDR-thresholded at $p < 0.05$.

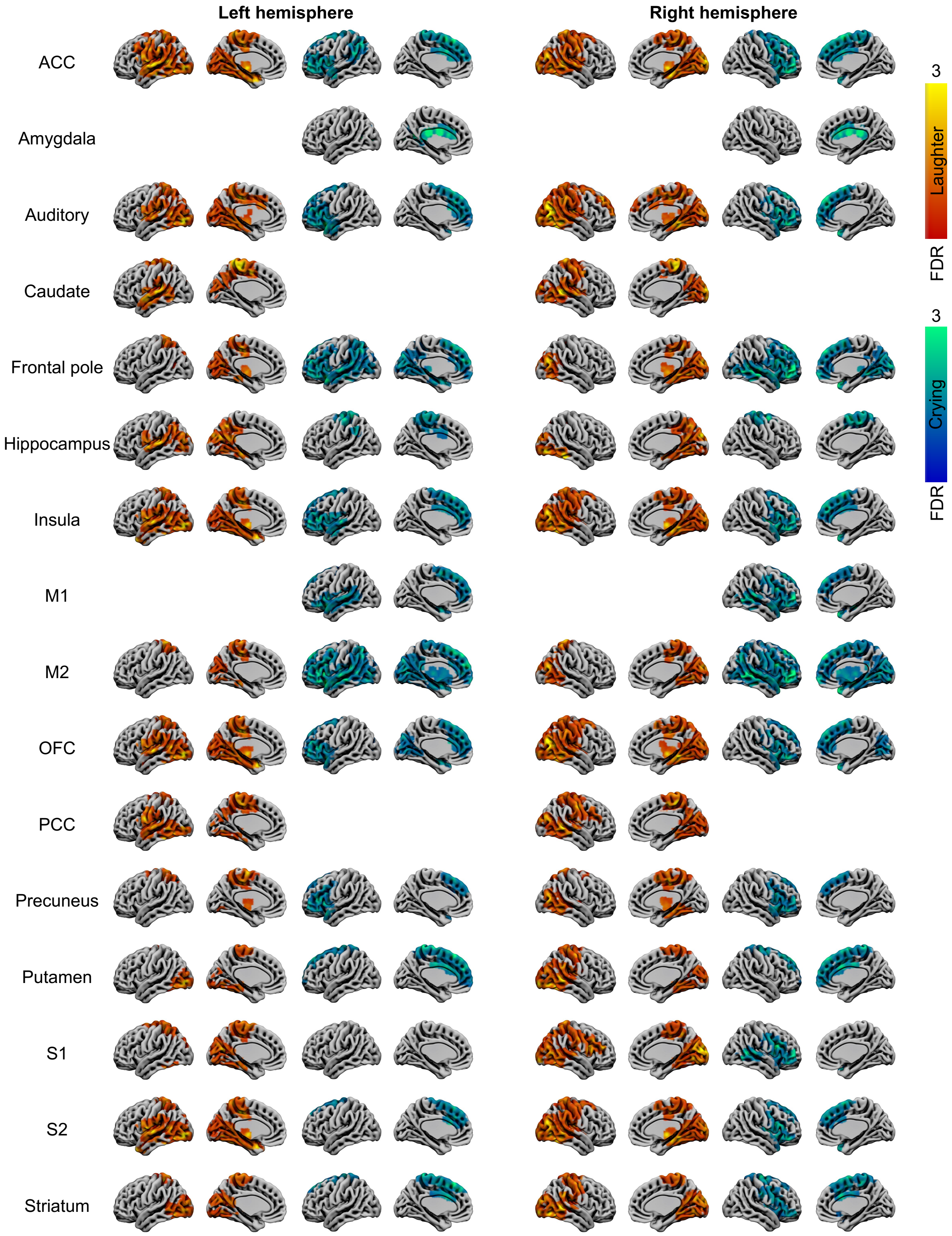


**Figure S2**. Negative correlations between regional MOR with BOLD response to laughter (hot colours) and crying (cool colours) sounds. The results are thresholded at $p < 0.05$ with FDR cluster level correction. ACC = anterior cingulate cortex, M1 = primary motor cortex, M2 = secondary motor cortex, OFC = orbitofrontal cortex, PCC = posterior cingulate cortex, S1 = primary somatosensory cortex, S2 = secondary somatosensory cortex.

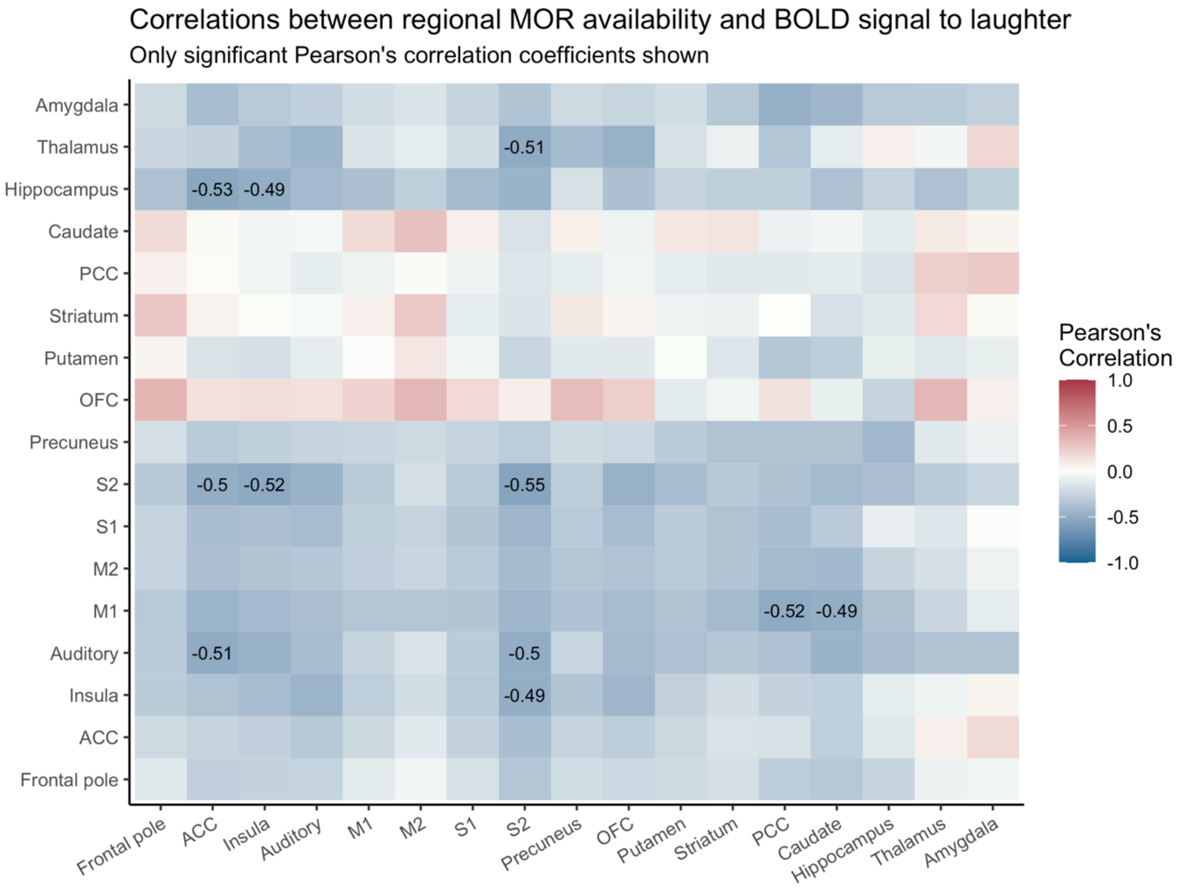


**Figure S3.** Correlation matrix for ROI-level MOR availability and ROI-level BOLD responses to laughter. ACC = anterior cingulate cortex, M1 = primary motor cortex, M2 = secondary motor cortex, OFC = orbitofrontal cortex, PCC = posterior cingulate cortex, S1 = primary somatosensory cortex, S2 = secondary somatosensory cortex.

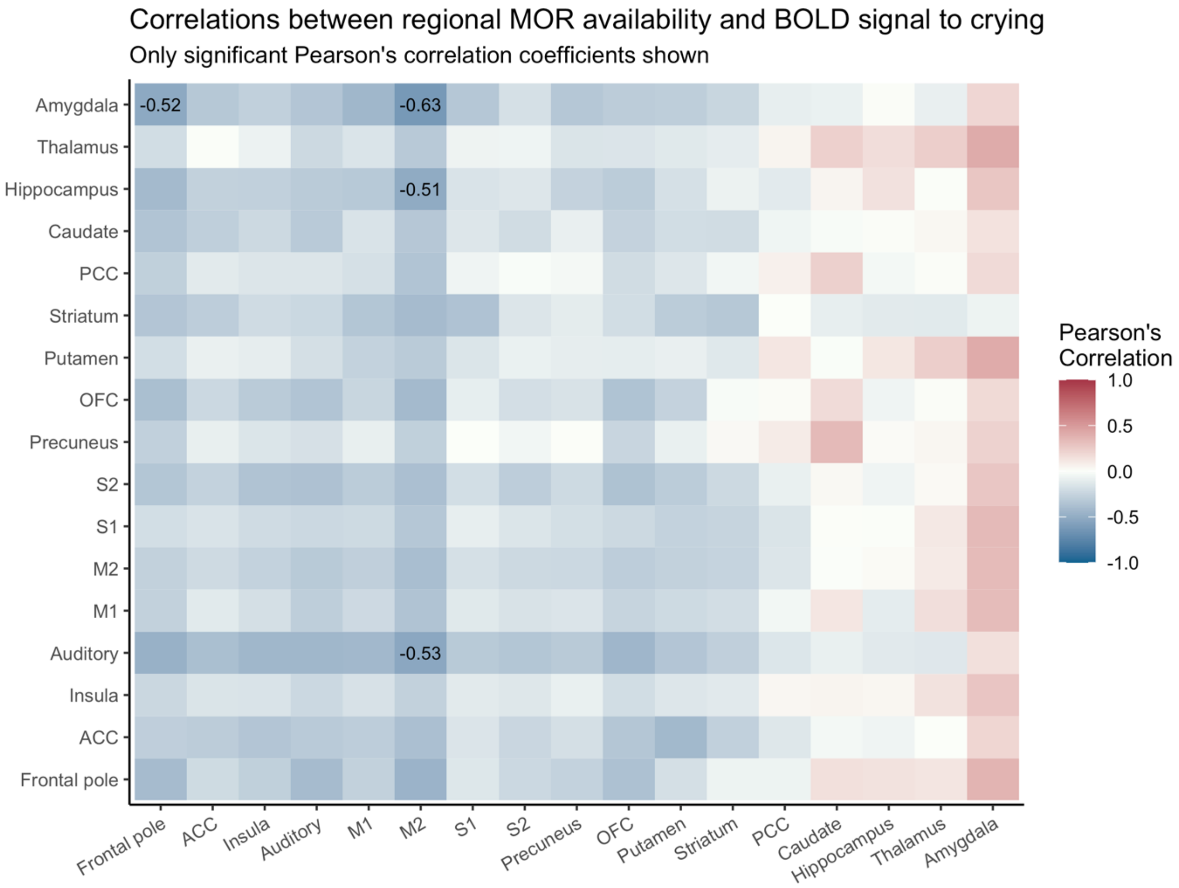


**Figure S4.** Correlation matrix for ROI-level MOR levels and ROI-level BOLD response to crying. ACC = anterior cingulate cortex, M1 = primary motor cortex, M2 = secondary motor cortex, OFC = orbitofrontal cortex, PCC = posterior cingulate cortex, S1 = primary somatosensory cortex, S2 = secondary somatosensory cortex.
